# Connectome-based predictive modeling shows sex differences in brain-based predictors of memory performance

**DOI:** 10.1101/2022.12.21.521314

**Authors:** Suyeon Ju, Corey Horien, Xilin Shen, Hamid Abuwarda, Anne Trainer, R Todd Constable, Carolyn A. Fredericks

**Author notes:** **Correspondence:** Carolyn A. Fredericks.

## Abstract

Alzheimer’s disease (AD) takes a more aggressive course in women than men, with higher prevalence and faster progression. Amnestic AD specifically targets the default mode network (DMN), which subserves short-term memory; past research shows relative hyperconnectivity in the posterior DMN in aging women. Higher reliance on this network during memory tasks may contribute to women’s elevated AD risk. Here, we applied connectome-based predictive modeling (CPM), a robust linear machine-learning approach, to the Lifespan Human Connectome Project-Aging (HCP-A) dataset (n=579). We sought to characterize sex-based predictors of memory performance in aging, with particular attention to the DMN. Models were evaluated using cross-validation both across the whole group and for each sex separately. Whole-group models predicted short-term memory performance with accuracies ranging from ρ=0.21-0.45. The best-performing models were derived from an associative memory task-based scan. Sex-specific models revealed significant differences in connectome-based predictors for men and women. DMN activity contributed more to predicted memory scores in women, while within- and between-visual network activity contributed more to predicted memory scores in men. While men showed more segregation of visual networks, women showed more segregation of the DMN. We demonstrate that women and men recruit different circuitry when performing memory tasks, with women relying more on intra-DMN activity and men relying more on visual circuitry. These findings are consistent with the hypothesis that women draw more heavily upon the DMN for recollective memory, potentially contributing to women’s elevated risk of AD.

## 1 Introduction

In addition to outnumbering men with Alzheimer’s disease (AD) by 2:1 (‘2022 Alzheimer’s disease facts and figures’, 2022), women with AD face faster accumulation of pathology and more severe illness with the same pathologic burden (Barnes *et al*., 2005; Buckley *et al*., 2018; Edwards *et al*., 2021). AD specifically targets the default mode network (DMN), which subserves short-term memory (Greicius *et al*., 2004; Sheline *et al*., 2010; Mormino *et al*., 2011; Brier *et al*., 2012). Yet, sex differences in the DMN over the course of aging, which may provide important clues to women’s higher vulnerability to AD, are poorly understood.

Prior research assessing sex differences in the aging brain has demonstrated that healthy aging women show lower segregation of functional networks (i.e., more cross-hemispheric/-module connections) (Ingalhalikar *et al*., 2014). Women have relatively higher DMN connectivity overall (Biswal *et al*., 2010; for the Women’s Brain Project and the Alzheimer Precision Medicine Initiative *et al*., 2018; Ritchie *et al*., 2018), and demonstrate higher connectivity than men in posterior DMN nodes, which relates to short-term memory performance (Ficek-Tani *et al*., In press).

Prediction-based approaches, in which models are built on training data and tested on unseen data, can help increase generalizability and reproducibility of findings (Yarkoni and Westfall, 2017; Scheinost *et al*., 2019; Poldrack, Huckins and Varoquaux, 2020; Marek *et al*., 2022; Yarkoni, 2022), and have the potential to generate useful biomarkers (Gabrieli, Ghosh and Whitfield-Gabrieli, 2015; Rosenberg, Casey and Holmes, 2018).

In this work, we use a predictive modeling-based approach to robustly characterize sex differences in the aging functional connectome. We used connectome-based predictive modeling (CPM) to predict short-term memory performance scores in a large dataset of healthy adults aged 36-100. We hypothesized that (a) predictive edges would vary substantially between men and women, (b) predictors would especially feature the DMN, with women relying more on within-DMN edges for memory task performance, and (c) women would show decreased network segregation than do men.

## 2. Methods

### 2.1 Participants

The data used were collected from participants enrolled in the Human Connectome Project-Aging (HCP-A) study (Bookheimer *et al*., 2019). Imaging data were from the 1.0 release of the HCP-A dataset, while the neurobehavioral data were from the 2.0 release. Imaging data consisted of 689 healthy subjects aged 36 to 100 from four data collection sites. See Bookheimer et al. (2019) for full exclusion criteria. As described previously (Ficek-Tani *et al*., In press), we implemented additional exclusion criteria based on motion (see below for details), missing data, and anatomical abnormalities. After exclusion, the remaining sample size was n=579 (330 female; 249 male).

Participants were well-matched in age, race, ethnicity, years of education, and handedness, but women outnumbered and outperformed men in global cognitive function (Montreal Cognitive Assessment), in-scanner memory task performance (FaceName task), and verbal learning (Rey Auditory Verbal Learning Test) (**Table 1**). Participants self-identified their sex at birth as male or female. While an “Other” option for sex was offered by the HCP-A study, no participants chose this option; gender identity was not assessed.

**Table 1.** Demographics and selected neuropsychological assessment and in-scanner task scores of HCP-A participants included in this study (Costa and McCrae, 1992; Nasreddine *et al*., 2005; Bean, 2011; Bookheimer *et al*., 2019). T-tests or chi square tests were performed as appropriate, excluding unknown/not reported values (Abbreviations: MOCA, Montreal Cognitive Assessment; RAVLT, Rey Auditory Verbal Learning Test).

|  | Female subjects | Male subjects | p-value |
| --- | --- | --- | --- |
| **Total number of participants** | 330 (57%) | 249 (43%) | N/A |
| **Age** | 56.94 (14.03) | 58.20 (14.13) | 0.284 |
| **Race** | American Indian/Alaska Native: 1<br>Asian: 19<br>Black/African American: 53<br>White: 231<br>More than one race: 18<br>Unknown/Not reported: 8 | American Indian/Alaska Native: 1<br>Asian: 26<br>Black/African American: 34<br>White: 177<br>More than one race: 9<br>Unknown/Not reported: 2 | 0.167 |
| **Ethnicity** | Hispanic or Latino: 36<br>Not Hispanic or Latino: 293<br>Unknown/Not reported: 1 | Hispanic or Latino: 18<br>Not Hispanic or Latino: 231<br>Unknown/Not reported: 0 | 0.216 |
| **Years of Education** | 15.38 (1.76) | 15.62 (1.82) | 0.121 |
| **Handedness** | Right: 283<br>Left: 21<br>Ambidextrous: 26 | Right: 197<br>Left: 21<br>Ambidextrous: 31 | 0.100 |
| **MOCA Total (points out of 30)** | 26.86 (2.24) | 26.21 (2.56) | 1.24E-3 |
| **FaceName Task Total (# face-name pairs recalled, out of 10)** | 6.97 (2.72) | 5.90 (2.96) | 1.27E-5 |
| **RAVLT Sum of Trials 1-5 (# words recalled, out of 75)** | 48.22 (9.71) | 44.06 (10.50) | 1.44E-6 |
| **RAVLT Trial 6 (# words recalled, out of 15)** | 10.13 (2.99) | 9.07 (3.31) | 7.47E-5 |

### 2.2 Imaging parameters

All subjects enrolled in HCP-A were scanned in a Siemens 3T Prisma scanner with 80mT/m gradients and 32-channel head coil. In addition to acquiring four resting-state fMRI (rfMRI) and three task-fMRI (tfMRI) scans per subject, structural MRI data (including one T1-weighted [T1w] scan) were also collected (Harms *et al*., 2018). In this study, we focus on the seven fMRI scans.

A multi-echo MPRAGE sequence (refer to (Harms *et al*., 2018) for scanning parameter details) was used for all T1w scans. A 2D multiband (MB) gradient-recalled echo (GRE) echo-planar imaging (EPI) sequence (MB8, TR/TE = 800/37 ms, flip angle = 52°) was used for all fMRI scans.

For each subject, four rfMRI scans consisting of 488 frames and lasting 6.5 minutes each (for a total of 26 minutes) were acquired, during which participants were instructed to remain awake while viewing a small white fixation cross in the center of a black background. The rfMRI scans were split between two sessions that occurred on the same day, with each session including one rfMRI with an anterior to posterior (AP) phase encoding direction and one rfMRI with a posterior to anterior (PA) direction.

The HCP-A includes the following three fMRI tasks, which were all programmed in PsychoPy (Peirce, 2007, 2008) and collected with PA phase encoding direction: Visuomotor (VisMotor), Conditioned Approach Response Inhibition Task (“CARIT” Go/NoGo task), and FaceName (Bookheimer *et al*., 2019). As below, we focus on the FaceName task scan both because of its relevance to short-term memory performance and because models derived from this scan outperform models derived from other scans. In the FaceName task, three blocks (encoding, distractor, and recall blocks) are repeated twice for each set of faces, totaling to a single, 276-second run. See (Harms *et al*., 2018) for full details on the HCP-A structural and functional MRI imaging parameters, and see (Bookheimer *et al*., 2019) for full details on tfMRI task administration.

### 2.3 Image preprocessing

The preprocessing approach has been described elsewhere (Greene *et al*., 2018; Horien *et al*., 2019). MPRAGE scans were skullstripped with optiBET (Lutkenhoff *et al*., 2014) and nonlinearly registered to the MNI template in BioImage Suite (BIS) (Joshi *et al*., 2011). BIS was used to linearly register each participant’s mean functional scan to their own MPRAGE scan. Participants were excluded from further analyses due to structural abnormalities after visually inspecting skullstripped and registered data. Functional data were motion-corrected using SPM8; participants whose scans showed maximum mean frame-to-frame displacement (FFD) above 0.3 mm were excluded to limit motion artifacts (Greene *et al*., 2018; Horien *et al*., 2018, 2019; Ju *et al*., 2020). Using Wilcoxon rank sum tests, we determined no differences in mean FFD between female and male subjects across all seven scan types (**Supplementary Table 1**). Linear, quadratic, and cubic drift, a 24-parameter model of motion (Satterthwaite *et al*., 2013), mean cerebrospinal fluid signal, mean white matter signal, and global signal were regressed from the data as described in (Ficek-Tani *et al*., In press).

### 2.4 Memory performance measures

Because we were interested in predictors of memory performance, we used performance on the FaceName task and the Rey Auditory Verbal Learning Test (RAVLT) as outcomes for our predictive models. For the FaceName task, participants were shown a total of 10 distinct faces, resulting in a maximum FaceName-Total Recall (FN-TR) score of 10 correctly identified faces. We also assessed both the learning (L) and immediate recall (IR) metrics from the RAVLT (Bean, 2011), a standard neuropsychological measure of declarative memory. In this assessment, a 15-word list is read to the participant, who is then asked to verbally recall as many as possible, five times. The total number of words recalled during this five-trial “learning period” sums to a RAVLT-L (“learning”) score out of 75 words. After being read a separate (interference) list and asked to recall it, the participant is read List A again, and the number of correctly-recalled words in this sixth trial is collected as the RAVLT-IR (“immediate recall”) score. RAVLT-IR is a sensitive metric for early-stage AD (Estévez-González *et al*., 2003).

### 2.5 Connectome-based predictive modeling

To predict memory performance using both rfMRI and tfMRI data from HCP-A, we used connectome-based predictive modeling (CPM), the details of which are described elsewhere (Shen *et al*., 2017).

In brief, connectivity matrices were constructed from each fMRI scan using the Shen 268-node atlas (Shen *et al*., 2013). These matrices and the memory performance scores of each participant were used to create our predictive models. Three subject groups were analyzed: all subjects, female-only, and male-only. Edges from connectivity matrices for each subject per scan were correlated to the three aforementioned memory performance measures, totaling to seven connectivity matrices and three memory scores per subject (21 total correlated matrices). Motion and age covariates were also included in the CPM analyses to account for in-scanner head motion, age, and their interaction in our predictions, as previously done (Scheinost *et al*., 2021; Dufford *et al*., 2022; Horien *et al*., 2022).

Using 5-fold cross validation, connectivity matrices and memory scores were divided into independent training (subjects from four of the folds) and testing (subjects in left-out fold) sets. Edge strength and memory were linearly related within the training set, and using a feature selection threshold of p = 0.01, a consensus connectivity matrix including only the edges most strongly positively or negatively correlated to memory was generated. Edge strengths in each subject’s connectivity matrix corresponding to the consensus matrix were summed into a single-subject connectivity value. A predictive model built using the linear relationship between the single-subject connectivity values and memory score was applied to the subjects in the testing set to generate memory performance predictions.

### 2.6 Model performance comparison

For all subject groups, Spearman’s correlation and root mean square error (defined as: 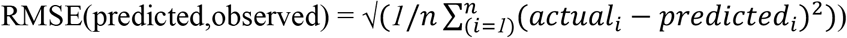were used to compare the similarity between predicted and observed memory scores to assess predictive model performance. After performing 1000 iterations of each CPM analysis, we selected the median-performing model to represent the model’s overall performance. To compare model performances between female and male groups for each fMRI scan, we used Wilcoxon rank sum tests.

We also tested our models against randomly permuted models by randomly shuffling participant labels prior to attempting to predict memory scores. After performing 1000 iterations of this permutation, we calculated the number of times the permuted predictive accuracy was greater than the median unpermuted prediction accuracy to generate a non-parametric p value, as done in (Scheinost *et al*., 2021):

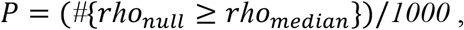

where #{*rho*_*null*_ ≥ *rho*_*median*_} indicates the number of permuted predictions numerically greater than or equal to the median of the unpermuted predictions. We applied the Benjamini-Hochberg procedure to these non-parametric p-values to control for multiple comparisons and correct for 21 tests for each of our three subject groups (Benjamini and Hochberg, 1995).

### 2.7 Inter-network significant-edge analyses

To visualize sex differences at the network level, we first split the aforementioned consensus matrix into two binarized matrices (a “positive” matrix containing edge with significant positive correlations to memory and the other “negative” matrix of edges with significant negative correlations to memory) for each predictive model. Categorization of nodes by functional network was determined using the 10-network parcellation of the Shen 268-node atlas (Horien *et al*., 2022). In this network grouping, the medial frontal (MF) network also includes some temporal and frontal nodes which often cluster with the DMN. Inter-network edges were defined as the number of significant edges between each pair of networks normalized by the total number of edges between the same network pair. As done in previous work, we defined edges as “significant” if they appear in at least 2 out of 5 folds in 40% of 1000 iterations of CPM to minimize noise while retaining meaningful connections (Rosenberg *et al*., 2016; Yip *et al*., 2019; Horien *et al*., 2022). In addition to using heatmaps to visualize the inter-network edges of both female and male groups separately, we subtracted male-group positive edges from female-group positive edges (and the same with the negative edges) across corresponding matrix cells to evaluate the inter-network sex differences.

### 2.8 Intra-network significant-edge analyses

Intra-network analyses were performed similarly to inter-network analyses above. Edges from binarized positively and negatively correlated connectivity matrices were summed across the 5 folds and 1000 iterations to generate a single value for each edge. These values were then used to generate the intra-DMN edge heatmap, with values ranging from -5000 (maximum negatively correlated) to 5000 (maximum positively correlated value). To evaluate differences in the “top-performing” nodes according to sex, individual edge values were summed across each row from the matrices and divided by 2 to account for the symmetric nature of the matrix, generating a summed vector (SV).

### 2.9 Network segregation analyses

We evaluated network segregation, a measure of the relative strength of within-network connections to between-network connections, using a novel association ratio metric. We defined the association ratio as the weighted sum of all edges within the network of interest, normalized by the weighted sum of all edges between this network and the whole set of regions of interest. Higher association ratio is therefore indicative of higher network segregation. To compare network segregation levels between sexes, we calculated and compared (using two-sample t-tests) the association ratio for certain networks of interest in women and men for each scan type.

Benjamini-Hochberg correction (see above) was applied to correct for 7 significance tests (for each model) across the 4 networks.

### 2.10 Data and code availability

Data from the HCP-A study are openly available (https://www.humanconnectome.org/study/hcp-lifespan-aging/data-releases). Image preprocessing was performed using BioImageSuite, a publicly-available software (https://medicine.yale.edu/bioimaging/suite/). Scripts for running CPM are available through GitHub (https://github.com/YaleMRRC/CPM). Other MATLAB scripts for CPM analyses can be found at https://github.com/frederickslab/CPM_HCP-A_sex_difference_study. Custom MATLAB colormap palettes were derived from ColorBrewer (http://colorbrewer.org/; Brewer, 2022).

## 3 Results

### 3.1 Model performance comparison

Please see **Supplementary Results** for details on model comparisons, including comparisons between models derived separately for each sex. Briefly, we trained and cross-validated models using functional connectivity data from all 7 scans to predict memory performance scores. Whole-group models robustly predicted all memory measures, with accuracies ranging from Spearman’s rho = 0.21 (RMSE = 3.34, p<0.0001) to rho = 0.45 (RMSE = 2.67, p<0.0001) across all models (**Supplementary Figure 2**). Models using the FaceName tfMRI scan consistently outperformed all other models; we therefore proceeded with models from this scan for the remaining analyses.

### 3.2 Inter-network significant-edge analyses

Visualizations of inter-network edges (number of significant edges normalized by network size) across all FaceName tfMRI models revealed differences in key edges predicting memory score for each sex. In particular, edges within the DMN and visual (visual I [VI], visual II [VII], and visual association areas [VAs]) networks showed the largest differences (**Figure 1, Supplementary Figure 6**). Given previous work showing measures of declarative verbal memory (including RAVLT metrics) can be predicted from the gray matter density of DMN structures, and because lower RAVLT-IR scores are associated with preclinical AD, we concentrated on the RAVLT-IR predictors derived from FaceName tfMRI models (Estévez-González *et al*., 2003; Moradi *et al*., 2017). In addition to visualizing the inter-network edges of females and males separately, we subtracted male-group edges from female-group edges across corresponding heatmap cells to evaluate inter-network differences between the sexes (**Figure 1**).

**Figure 1.**
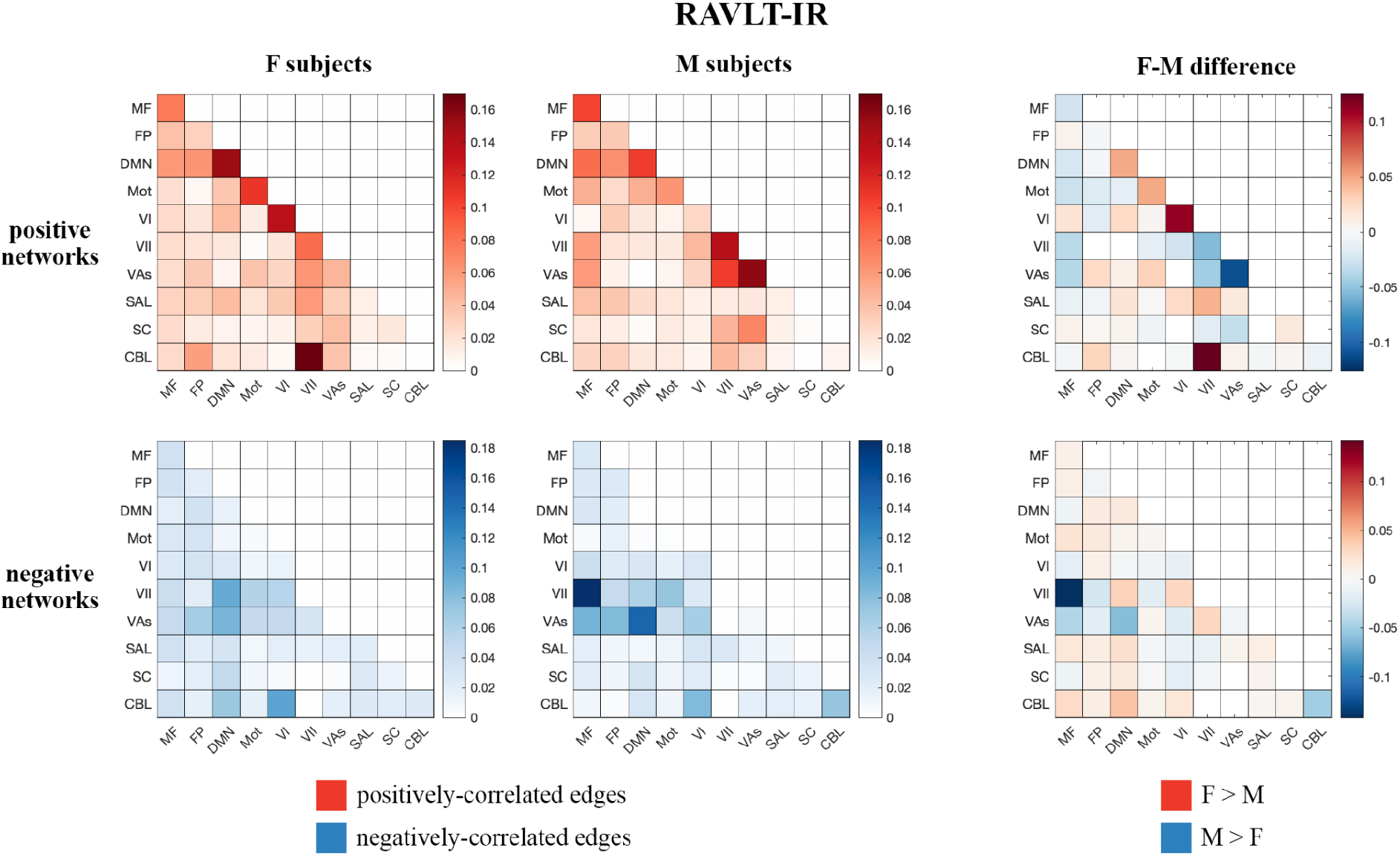
Positive and negative matrices from the RAVLT-IR-predicting model showing inter-network connections (number of significant edges normalized by network size for each network pair) for female and male subjects, as well as the difference between both sexes (derived by subtracting male inter-network edges from female inter-network edges). Both sexes show positive predictors in the intra-DMN edges. Female subjects show more positive predictors in the intra-VI-network edges relative to male subjects, while male subjects show more positive predictors in the intra- and inter-visual (VII and VAs)-network edges relative to female subjects. Negative predictors of both sexes relied on edges between DMN and visual networks; however, male subjects’ negative predictors relied more on edges between the MF and VII networks than those of female subjects (Abbreviations: F, female; M, male; MF, medial frontal; FP, fronto-parietal; DMN, default mode network; Mot, motor; VI, visual I; VII, visual II; VAs, visual association areas; SAL, limbic; SC, basal ganglia; CBL, cerebellum; RAVLT-IR, RAVLT-Immediate Recall).

Both sexes show positive predictors with intra-DMN edges, with female scores predicting intra-DMN connectivity more strongly than those of males. Female positive predictors also relied more strongly on intra-VI edges than those of males, while male positive predictors relied more strongly on the intra- and inter-network connectivity of the VII and VAs networks relative to those of females. Both sexes displayed negative predictors with edges between DMN and visual networks; however, males show more negative predictors with edges between the MF and VII networks, as well as between the DMN and VII networks, relative to females.

### 3.3 Intra-network significant-edge analyses

Given the preferential contribution of intra-DMN edges to the female models, we examined all intra-DMN edges and evaluated their strengths in male and female models. To do so, we generated a heatmap of intra-DMN edges (**Figure 2**). In the RAVLT-IR model, we found that edges from more posterior DMN nodes were preferentially increased in females as opposed to males. This trend held true for the RAVLT-L model and FN-TR models (**Supplementary Figure 7**). Negatively correlated edges negligibly contributed to both male and female models (**Figure 2, Supplementary Figure 7**). Both sexes displayed strong connections in the left posterior cingulate cortex (L PCC) and precuneus, known hubs of the DMN.

**Figure 2.**
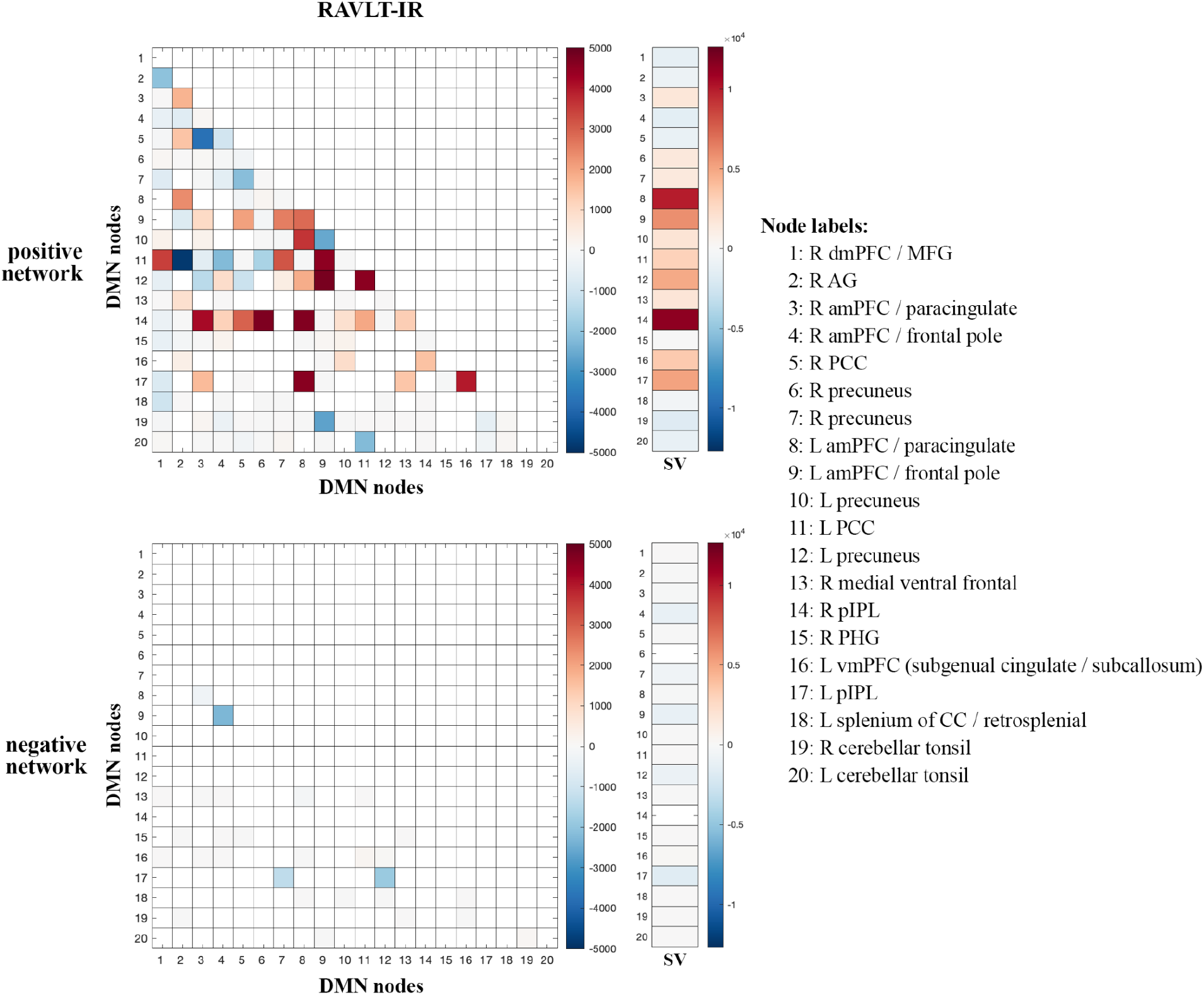
Intra-DMN connectivity differences between males and females. Intra-DMN edge counts from the RAVLT-IR models were calculated and plotted as a heat map (female – male edge counts). Red indicates higher female counts and blue indicates higher male counts for each edge (Abbreviations: RAVLT-IR, RAVLT-Immediate Recall; L, left; R, right; dmPFC, dorsomedial prefrontal cortex; MFG, middle frontal gyrus; AG, angular gyrus; aMPFC, anterior medial prefrontal cortex; PCC, posterior cingulate cortex; pIPL, posterior inferior parietal lobe; PHG, parahippocampal gyrus; vmPFC, ventromedial prefrontal cortex; CC, corpus callosum; SV, summed vector).

To summarize node-level differences, we summed the number of edges associated with each node and found consistent female preference for activity of the right posterior inferior parietal lobe (R pIPL) and left anterior medial prefrontal cortex (L amPFC)/paracingulate cortex (**Figure 2**). The R pIPL was consistently and preferentially elevated in all female models analyzed (**Supplementary Figure 7**). This analysis demonstrates differential edge- and node-level contributions to male and female models.

### 3.4 Network segregation analyses

We then evaluated and compared a metric of network segregation (see Methods, “Network Segregation Analysis”) within the DMN and visual (VI, VII, VAs) networks between females and males, given the strong brain-behavior correlations in these networks across all memory performance outcomes. Our analysis demonstrated increased network segregation of the DMN in females relative to males, and increased network segregation of VII and VAs in males relative to females (**Table 2**). Additionally, these findings echoed our previous CPM analysis results in that we also observed sex differences in neurobiological organization.

**Table 2.** Network segregation differences between female and male subjects. Two-sample t-tests comparing the association ratios for networks of interest between the sexes revealed increased DMN segregation in female subjects and increased VII and VAs network segregation in male subjects. Red indicates significantly higher network segregation in female subjects than male subjects and blue indicates significantly higher network segregation in male subjects than female subjects. We report these results as ‘t-statistic (p-value)’ in the table. † indicates the models that did not survive correction for multiple comparisons.

| Scan Type | Default Mode Network (DMN) | Visual I (VI) Network | Visual II (VII) Network | Visual Association Areas (VAs) |
| --- | --- | --- | --- | --- |
| REST1_AP | 3.17 (0.0016) | -1.32 (0.1879)† | -9.02 (2.69E-18) | -4.32 (1.87E-05) |
| REST1_PA | 3.45 (0.0006) | -0.40 (0.6920)† | -7.79 (3.18E-14) | -3.92 (0.0001) |
| REST2_AP | 1.21 (0.2259)† | -0.92 (0.3557)† | -7.07 (4.42E-12) | -5.16 (3.47E-07) |
| REST2_PA | 2.04 (0.0419)† | 0.07 (0.9425)† | -6.38 (3.66E-10) | -2.55 (0.0111) |
| CARIT | 2.57 (0.0104) | 2.11 (0.0349)† | -5.33 (1.40E-07) | -0.54 (0.5864)† |
| FACENAME | 1.18 (0.2397)† | 1.74 (0.0821)† | -4.46 (9.71E-06) | -1.03 (0.3017)† |
| VISMOTOR | 0.20 (0.8399)† | -0.47 (0.6360)† | -3.33 (0.0009) | -3.06 (0.0023) |

## 4 Discussion

We use CPM to identify sex differences in the functional connectivity underlying memory performance in a large sample of healthy aging adults. We provide evidence that distinct edges for men and women predict short-term verbal memory task performance, and that within-DMN edges contribute more to memory scores in females than in males. Predictive edges for males, in contrast, include more edges within and across visual sensory and association networks. In contrast to prior literature suggesting globally decreased network segregation in older women compared with men, we also show higher segregation of the DMN (but lower segregation of visual sensory and association networks) in women.

These findings imply that when compared with males, females have a higher reliance upon connections within the DMN, the intrinsic connectivity network targeted in AD, in performing memory-related tasks. Increased DMN connectivity, particularly in posterior nodes, has been associated with vulnerability to Alzheimer’s disease (Bookheimer *et al*., 2000; Filippini *et al*., 2009; Sperling *et al*., 2009; Mormino *et al*., 2011; Schultz *et al*., 2017); increased connectivity in preclinical AD settings is thought to represent the compensatory response of a network under stress (Bondi *et al*., 2005; Filippini *et al*., 2009; Qi *et al*., 2010; Mormino *et al*., 2011), and symptomatic disease is associated with progressive hypoconnectivity across the network (Greicius *et al*., 2004; Sheline *et al*., 2010; Brier *et al*., 2012).

This study and our previous findings in the same dataset (Ficek-Tani *et al*., In press) converge on an emerging narrative of increased connectivity and functional segregation of the DMN in aging women. Women rely upon specific DMN edges for memory performance; connections between the bilateral pIPL and the two greatest hubs of the DMN, the mPFC and the PCC/precuneus are the strongest predictors. Our prior work suggests that women have relatively increased within-DMN connectivity compared with men, particularly in posterior nodes and particularly during perimenopausal decades (Ficek-Tani *et al*., In press). Reliance upon intra-DMN edges for memory performance likely has its advantages: we and others have shown that DMN connectivity, particularly between posterior nodes, correlates with memory task performance (Fredericks *et al*., 2019; Natu *et al*., 2019; Kang *et al*., 2021; Vanneste *et al*., 2021; Ficek-Tani *et al*., In press), and the literature consistently demonstrates that women outperform men across the lifespan in tests of verbal episodic memory (Bleecker *et al*., 1988; Herlitz, Nilsson and Bäckman, 1997; Golchert *et al*., 2019).

We also find relatively greater functional segregation of the DMN in women than in men. Functional segregation (i.e., reliance on within-more than between-network connectivity to perform a network-associated task) declines across the brain with aging, and is associated with decreased performance on tests of attention and memory performance (Chan *et al*., 2014; Geerligs *et al*., 2015; Ng *et al*., 2016). AD pathology is associated with decreased functional segregation (Cassady *et al*., 2021), and prior work in this field has suggested that women show decreased functional segregation over the course of aging and during memory task performance specifically (Ingalhalikar *et al*., 2014; Rabipour *et al*., 2021; Subramaniapillai *et al*., 2022), potentially relating to AD vulnerability (Rabipour *et al*., 2021). We show that sex differences in segregation are network-specific: women have relatively decreased segregation of visual sensory and visual association networks, but increased DMN segregation relative to men.

## 5 Limitations and Future Directions

While the HCP-A dataset has many strengths, it has limitations. Specifically, while the dataset is large and offers very high-quality neuroimaging and neuropsychological characterization, it is cross-sectional, so we cannot assess for longitudinal effects. Second, amyloid biomarkers are not available for the participants, so we cannot examine the effect of preclinical AD on the measures of interest.

In terms of our results, we identify specific edges within the brain connectome and within the DMN in particular that contribute to memory performance in women specifically. The translational impact of these findings will depend on future work investigating whether these edges share a common gene expression pattern or other characteristic at the cellular level, which could be leveraged towards a potential therapeutic target. Additionally, our analyses suggest that edges between the visual sensory networks and the cerebellum may play an important role in memory performance, particularly for women. Future analyses that parcellate the cerebellum will be important for interpreting this finding, given that the cerebellum participates in many intrinsic connectivity networks (Buckner *et al*., 2011).

Finally, our work addresses the impact of self-reported sex on network changes, but AD risk in women also depends upon gender-based factors such as lack of access to activities which promote cognitive reserve, such as cardiovascular exercise, occupational complexity, and educational attainment (Mielke, Vemuri and Rocca, 2014). Additionally, the interplay of assigned sex at birth and gender identity was not assessed due to a lack of the required information in the HCP dataset. While we used self-identified sex to distinguish subjects, this categorization may not capture the complex dynamics that may contribute to the sex differences described above. Future work should seek to incorporate other variables, as has been recently suggested regarding ovarian hormone status (Rocks, Cham and Kundakovic, 2022), and to incorporate metrics of cognitive reserve.

## 6 Conclusion

In summary, this study makes three key contributions to our understanding of sex differences in brain circuitry driving memory performance, which could have implications for women’s higher vulnerability to AD. First, we found that women relied more on within-network DMN edges (specifically bilateral posterior inferior parietal lobe and its connections to the major DMN hubs, medial prefrontal cortex and posterior cingulate/precuneus) for memory task performance than did men. Second, we determined that men’s memory task performance was predicted by edges distributed more broadly both within and between visual sensory and visual association networks and the medial frontal network. Finally, in contrast to prior literature which suggests increased generalization of cognitive circuits in aging women, we show that women have relatively greater functional segregation of the DMN than men during memory task performance.

This work adds to the growing literature suggesting that women rely more on the DMN than do men both at rest and during memory task performance. At rest, women have relatively higher DMN connectivity (Biswal *et al*., 2010; Scheinost *et al*., 2015; Cavedo *et al*., 2018; Ritchie *et al*., 2018; Ficek-Tani *et al*., In press), with higher posterior DMN connectivity particularly during the menopausal decades (Ficek-Tani *et al*., In press); this increased connectivity correlates with better performance on tests of short-term memory (Fredericks *et al*., 2019; Natu *et al*., 2019; Kang *et al*., 2021; Vanneste *et al*., 2021; Ficek-Tani *et al*., In press). This profile is similar to individuals with preclinical (amyloid-β +) or elevated genetic risk (e.g. APOE-ε4+) for AD (Bookheimer *et al*., 2000; Filippini *et al*., 2009; Sperling *et al*., 2009; Mormino *et al*., 2011; Schultz *et al*., 2017).

We need to understand why AD has a more aggressive phenotype in women. Taken together this work adds to a body of literature that suggests that women’s relative increased reliance on within-DMN connectivity could lead to “overuse” and vulnerability of this network to pathology over time. Future work examining the common cellular features of the nodes composing women’s strongest predictive edges have the potential to translate as therapeutic targets.

## 8 Author Contributions

SJ, CH, RTC, and CF conceived of the presented idea. SJ and CF developed the theory and specified assessment parameters. CH, XS, and RTC developed computational models. SJ, CH, HA, and AT performed computations. SJ performed visualizations of the results. RTC helped supervise the project. CF supervised the project. SJ, CH, HA, AT, and CF drafted the original manuscript. All authors discussed the results and contributed to the manuscript.

### 9 Funding

This work was funded by awards to Dr. Fredericks from the National Institutes of Health (5K23AG059919-04), the Alzheimer’s Association (2019-AACSF-644153), Women’s Health Research at Yale, and the McCance Foundation. Research reported in this publication was supported by the National Institute on Aging of the National Institutes of Health under Award Number U01AG052564 and by funds provided by the McDonnell Center for Systems Neuroscience at Washington University in St. Louis. The content is solely the responsibility of the authors and does not necessarily represent the official views of the National Institutes of Health.

## 10. Acknowledgements

We thank Bronte Ficek-Tani for her assistance in preprocessing the HCP-A fMRI data used in this study and the Human Connectome Project - Lifespan team for making the HCP-A dataset available to the research community. We also thank the National Institutes of Health (5K23AG059919-04), the Alzheimer’s Association (2019-AACSF-644153), Women’s Health Research at Yale, and the McCance Foundation for their support of this work.

