## Supplementary Materials for "Connectome-based predictive modeling shows sex differences in brain-based predictors of memory performance"

Supplementary Material

**1	Supplementary Methods**

Using CPM, we trained and cross-validated models using functional connectivity data from all seven scans to predict memory performance scores. Whole-group models robustly predicted all memory measures, with accuracies ranging from Spearman’s rho = 0.21 (RMSE = 3.34, p < 0.0001) to rho = 0.45 (RMSE = 2.67, p < 0.0001) across all models (**Supplementary Figure 1**); see **Supplementary Table 2** for exact rho and significance values. Specifically, the models derived using the FaceName tfMRI scan consistently outperformed all other models; Steigler’s Z tests were performed between the FaceName model and all other models within each memory measure to confirm this (**Supplementary Table 3**; Weiss, 2011).

We also derived predictive models separately for both female and male subjects, the majority of which also robustly predicted all memory measures (the only exceptions being the three male-group models indicated in **Supplementary Table 5**). While the female and male models generated with the FaceName task scan still outperformed models generated with all other fMRI scans within each subject group, the FaceName tfMRI was the only one across all memory scores in which male models outperformed female ones (**Supplementary Figure 2**). The one exception to this is the RAVLT-L-predicting model generated with the REST1_PA resting scan, though we suspect that this scan is an outlier amongst the rfMRIs (as the female-male model performance patterns for the other three rfMRIs are more consistent). See **Supplementary Tables 4 and 5** for exact rho and significance values. Wilcoxon rank sum tests revealed that female- and male-group model performances differed across all fMRI scans for each memory score (indicated by asterisks in **Supplementary Figure 2**; see test statistics in **Supplementary Table 6**). All scans across the three subject groups except the REST1_AP - FN-TR-predicting, REST1_PA - FN-TR-predicting, and VisMotor - RAVLT-L-predicting male-group models (indicated in **Supplementary Table 5**) were significant after multiple comparisons correction.

Across all subject group models, the FaceName tfMRI scan models generated the strongest predictors of all memory performance measures. For this reason, we focused all further analyses in this study on the FaceName tfMRI models.

**2	Supplementary Data**

|  | **F subjects (n=330)** | | **M subjects (n=249)** | | **F+M subjects** | **F vs. M t-test** |
| --- | --- | --- | --- | --- | --- | --- |
| **Scan Type** | **Mean FFD** | **Standard Deviation** | **Mean FFD** | **Standard Deviation** | **Mean FFD** | **p-value** |
| REST1_AP | 0.1251 | 0.0557 | 0.1168 | 0.0494 | 0.1216 | 0.1513 |
| REST1_PA | 0.1123 | 0.0481 | 0.1079 | 0.0455 | 0.1104 | 0.3164 |
| REST2_AP | 0.1318 | 0.0595 | 0.122 | 0.0517 | 0.1276 | 0.1121 |
| REST2_PA | 0.1176 | 0.0521 | 0.1092 | 0.0455 | 0.114 | 0.1047 |
| CARIT | 0.1121 | 0.0458 | 0.1078 | 0.0426 | 0.1102 | 0.3418 |
| FACENAME | 0.1161 | 0.0463 | 0.111 | 0.0441 | 0.1139 | 0.1975 |
| VISMOTOR | 0.1157 | 0.0491 | 0.1105 | 0.0438 | 0.1135 | 0.3445 |

**Supplementary Table 1.** Motion (mean FFD) data for each scan type in the HCP-A n=579 cohort. No motion differences were detected between female and male subjects across all scan types (p-values determined using Wilcoxon rank sum tests; abbreviations: F, female; M, male; n, number of subjects; FFD, frame-to-frame displacement).

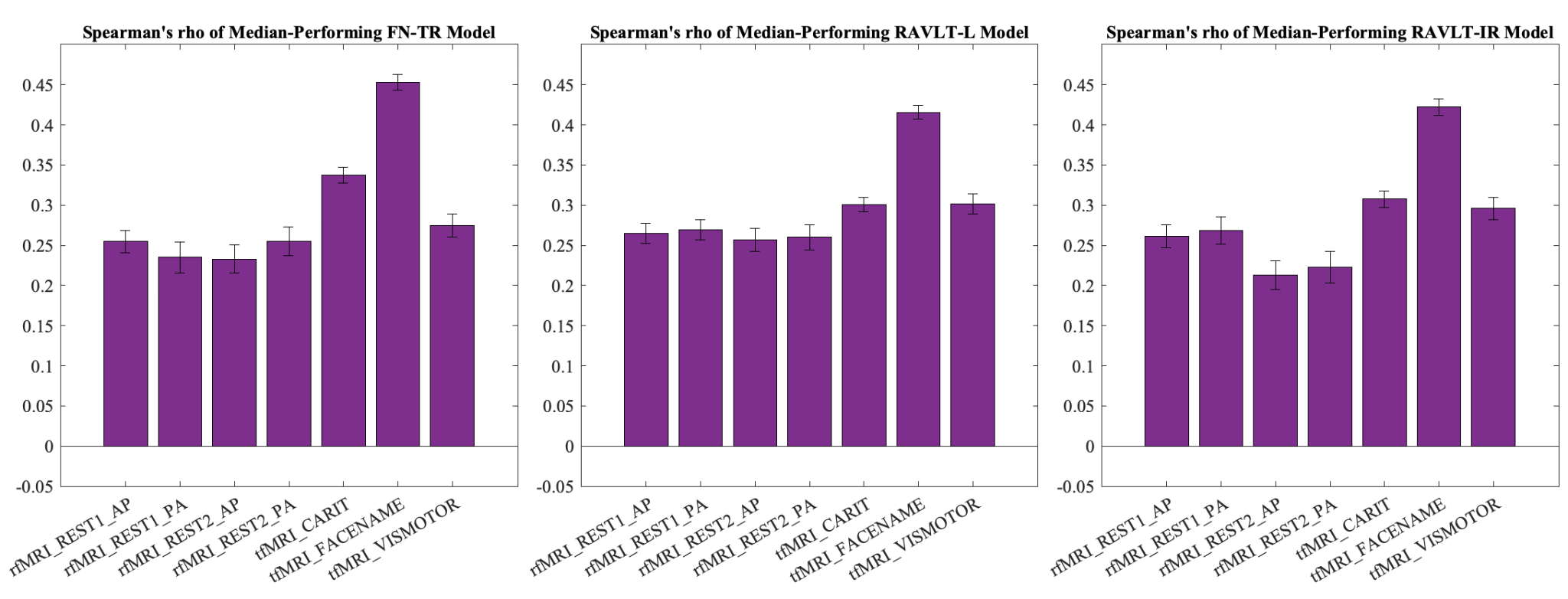

**Supplementary Figure 1**. Whole-group predictive model performance (measured using Spearman’s correlation coefficient (rho) values of the median-performing iterations of each model) across all fMRI scans for each memory metric (FN-TR, RAVLT-L, RAVLT-IR). The model derived from the FaceName task scan consistently outperformed other models across all memory scores. Error bars represent standard deviations of the 1000 iterations of each model (Abbreviations: rfMRI, resting-state fMRI; tfMRI, task fMRI; FN-TR, FaceName-Total Recall; RAVLT, Rey Auditory Verbal Learning Test; RAVLT-L, RAVLT-Learning; RAVLT-IR, RAVLT-Immediate Recall).

| **FN-TR-predicting models** | | | | |
| --- | --- | --- | --- | --- |
| **Scan Type** | **Spearman's rho** | **Spearman's p-value** | **RMSE** | **Permutation testing p-value** |
| REST1_AP | 0.2547 | 9.99E-10 | 2.9091 | <0.0001 |
| REST1_PA | 0.2351 | 1.85E-08 | 2.9465 | <0.0001 |
| REST2_AP | 0.2330 | 2.49E-08 | 2.9586 | <0.0001 |
| REST2_PA | 0.2550 | 9.49E-10 | 2.9389 | <0.0001 |
| CARIT | 0.3372 | 2.49E-16 | 2.8225 | <0.0001 |
| FACENAME | 0.4532 | 1.15E-29 | 2.6658 | <0.0001 |
| VISMOTOR | 0.2750 | 3.73E-11 | 2.8849 | <0.0001 |
| **RAVLT-L-predicting models** | | | | |
| **Scan Type** | **Spearman's rho** | **Spearman's p-value** | **RMSE** | **Permutation testing p-value** |
| REST1_AP | 0.2648 | 1.54E-10 | 10.2880 | <0.0001 |
| REST1_PA | 0.2694 | 7.17E-11 | 10.3227 | <0.0001 |
| REST2_AP | 0.2570 | 5.49E-10 | 10.4488 | <0.0001 |
| REST2_PA | 0.2602 | 3.29E-10 | 10.3883 | <0.0001 |
| CARIT | 0.3010 | 2.59E-13 | 10.1519 | <0.0001 |
| FACENAME | 0.4156 | 4.78E-25 | 9.7426 | <0.0001 |
| VISMOTOR | 0.3020 | 2.13E-13 | 10.1907 | <0.0001 |
| **RAVLT-IR-predicting models** | | | | |
| **Scan Type** | **Spearman's rho** | **Spearman's p-value** | **RMSE** | **Permutation testing p-value** |
| REST1_AP | 0.2611 | 2.11E-10 | 3.2404 | <0.0001 |
| REST1_PA | 0.2685 | 6.18E-11 | 3.2409 | <0.0001 |
| REST2_AP | 0.2130 | 2.58E-07 | 3.3388 | <0.0001 |
| REST2_PA | 0.2230 | 6.67E-08 | 3.3063 | <0.0001 |
| CARIT | 0.3076 | 4.82E-14 | 3.1918 | <0.0001 |
| FACENAME | 0.4222 | 3.25E-26 | 3.0289 | <0.0001 |
| VISMOTOR | 0.2961 | 4.44E-13 | 3.2289 | <0.0001 |

**Supplementary Table 2.** Whole-group predictive model performance results of the median-performing iteration of 1000 total iterations per model. All scans were significant after multiple comparisons correction (Abbreviations: FN-TR, FaceName-Total Recall; RAVLT, Rey Auditory Verbal Learning Test; RAVLT-L, RAVLT-Learning; RAVLT-IR, RAVLT-Immediate Recall; RMSE, root-mean-square error).

| **Scan being compared to FACENAME scan** | **FN-TR-predicting models** | **RAVLT-L-predicting models** | **RAVLT-IR-predicting models** |
| --- | --- | --- | --- |
| REST1_AP | <0.001 | <0.001 | <0.001 |
| REST1_PA | <0.001 | <0.001 | <0.001 |
| REST2_AP | <0.001 | <0.001 | <0.001 |
| REST2_PA | <0.001 | <0.001 | <0.001 |
| CARIT | <0.001 | <0.001 | <0.001 |
| VISMOTOR | <0.001 | <0.001 | <0.001 |

**Supplementary Table 3:** P-values from Steigler’s Z tests between the FaceName model and all other models within each memory metric. These results show that the FaceName tfMRI scan consistently outperformed all other models across all memory scores (Abbreviations: FN-TR, FaceName-Total Recall; RAVLT, Rey Auditory Verbal Learning Test; RAVLT-L, RAVLT-Learning; RAVLT-IR, RAVLT-Immediate Recall; RMSE, root-mean-square error).

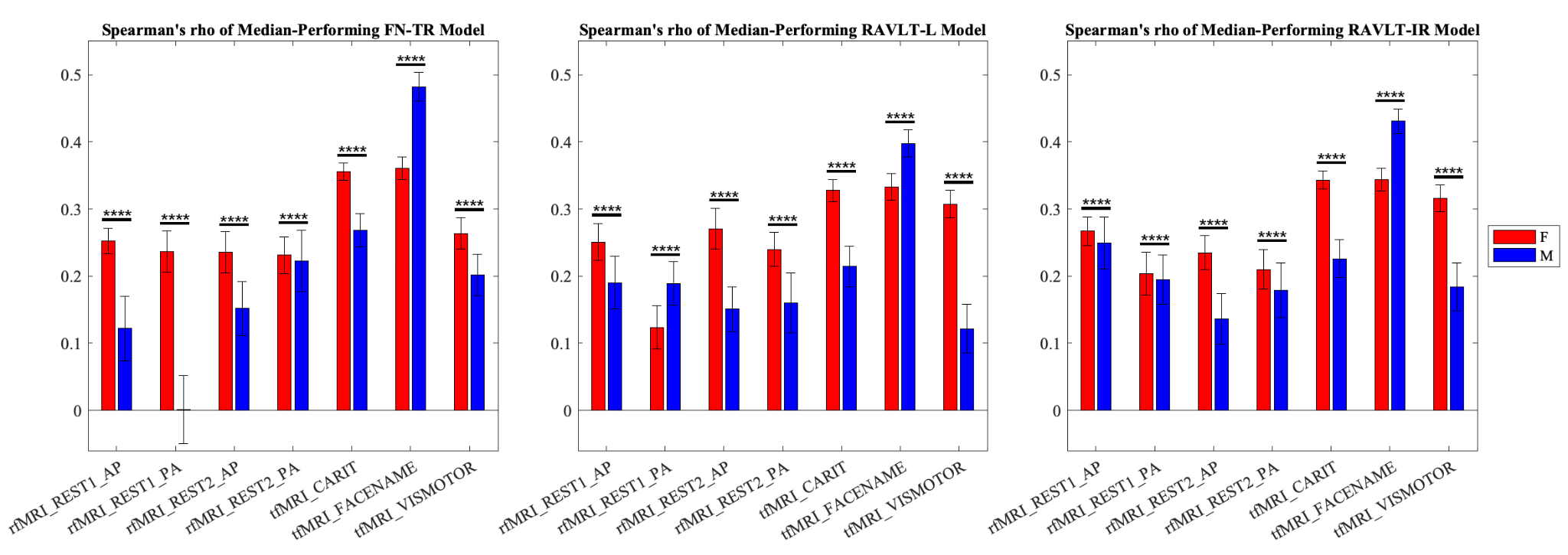

**Supplementary Figure 2.** Sex differences in predictive model performance across all fMRI scans for each memory metric. Models derived from female participants outperform models derived from males in most fMRI scan models, while models derived from male participants perform best in models derived from the FaceName tfMRI scan. Red indicates rho values of median-performing female-group models, and blue indicates rho values of median-performing male-group models. Error bars represent standard deviations of the 1000 iterations of each model (* = p < 0.05, ** = p < 0.01, *** = p < 0.001, **** = p < 0.0001; abbreviations: F, female; M, male; rfMRI, resting-state fMRI; tfMRI, task fMRI; FN-TR, FaceName-Total Recall; RAVLT, Rey Auditory Verbal Learning Test; RAVLT-L, RAVLT-Learning; RAVLT-IR, RAVLT-Immediate Recall).

| **FN-TR-predicting models** | | | | |
| --- | --- | --- | --- | --- |
| **Scan Type** | **Spearman's rho** | **Spearman's p-value** | **RMSE** | **Permutation testing**  **p-value** |
| REST1_AP | 0.2521 | 4.95E-06 | 2.7355 | <0.0001 |
| REST1_PA | 0.2361 | 1.97E-05 | 2.8031 | <0.0001 |
| REST2_AP | 0.2357 | 2.05E-05 | 2.7569 | 0.001 |
| REST2_PA | 0.2313 | 2.94E-05 | 2.7860 | 0.001 |
| CARIT | 0.3557 | 5.61E-11 | 2.6529 | <0.0001 |
| FACENAME | 0.3606 | 2.92E-11 | 2.6060 | <0.0001 |
| VISMOTOR | 0.2637 | 1.73E-06 | 2.7358 | <0.0001 |
| **RAVLT-L-predicting models** | | | | |
| **Scan Type** | **Spearman's rho** | **Spearman's p-value** | **RMSE** | **Permutation testing**  **p-value** |
| REST1_AP | 0.2507 | 5.07E-06 | 10.7916 | <0.0001 |
| REST1_PA | 0.1237 | 2.63E-02 | 11.3774 | 0.023 |
| REST2_AP | 0.2706 | 7.90E-07 | 10.6690 | <0.0001 |
| REST2_PA | 0.2400 | 1.30E-05 | 10.8768 | <0.0001 |
| CARIT | 0.3277 | 1.59E-09 | 10.5106 | <0.0001 |
| FACENAME | 0.3328 | 8.63E-10 | 10.5514 | <0.0001 |
| VISMOTOR | 0.3075 | 1.68E-08 | 10.6258 | <0.0001 |
| **RAVLT-IR-predicting models** | | | | |
| **Scan Type** | **Spearman's rho** | **Spearman's p-value** | **RMSE** | **Permutation testing**  **p-value** |
| REST1_AP | 0.2672 | 9.46E-07 | 3.0664 | <0.0001 |
| REST1_PA | 0.2038 | 2.08E-04 | 3.1409 | <0.0001 |
| REST2_AP | 0.2349 | 1.78E-05 | 3.0945 | <0.0001 |
| REST2_PA | 0.2100 | 1.30E-04 | 3.1348 | <0.0001 |
| CARIT | 0.3431 | 1.83E-10 | 2.9764 | <0.0001 |
| FACENAME | 0.3439 | 1.65E-10 | 2.9655 | <0.0001 |
| VISMOTOR | 0.3160 | 5.14E-09 | 3.0390 | <0.0001 |

**Supplementary Table 4.** Female-group predictive model performance results of the median-performing iteration of 1000 total iterations per model. All scans were significant after multiple comparisons correction (Abbreviations: FN-TR, FaceName-Total Recall; RAVLT, Rey Auditory Verbal Learning Test; RAVLT-L, RAVLT-Learning; RAVLT-IR, RAVLT-Immediate Recall; RMSE, root-mean-square error).

| **FN-TR-predicting models** | | | | |
| --- | --- | --- | --- | --- |
| **Scan Type** | **Spearman's rho** | **Spearman's p-value** | **RMSE** | **Permutation testing p-value** |
| REST1_AP | 0.1221 | 5.96E-02 | 3.2437 | 0.061 † |
| REST1_PA | 0.0013 | 9.84E-01 | 3.3472 | 0.457 † |
| REST2_AP | 0.1516 | 1.90E-02 | 3.1779 | 0.03 |
| REST2_PA | 0.2230 | 5.14E-04 | 3.0916 | 0.003 |
| CARIT | 0.2684 | 2.62E-05 | 3.0442 | <0.0001 |
| FACENAME | 0.4821 | 2.57E-15 | 2.7930 | <0.0001 |
| VISMOTOR | 0.2015 | 1.74E-03 | 3.0970 | 0.004 |
| **RAVLT-L-predicting models** | | | | |
| **Scan Type** | **Spearman's rho** | **Spearman's p-value** | **RMSE** | **Permutation testing p-value** |
| REST1_AP | 0.1901 | 2.93E-03 | 11.7466 | 0.006 |
| REST1_PA | 0.1893 | 3.05E-03 | 11.7947 | 0.006 |
| REST2_AP | 0.1509 | 1.86E-02 | 12.1197 | 0.025 |
| REST2_PA | 0.1601 | 1.25E-02 | 11.9080 | 0.027 |
| CARIT | 0.2146 | 7.59E-04 | 11.5681 | 0.001 |
| FACENAME | 0.3979 | 1.20E-10 | 10.8492 | <0.0001 |
| VISMOTOR | 0.1215 | 5.86E-02 | 12.0702 | 0.053 † |
| **RAVLT-IR-predicting models** | | | | |
| **Scan Type** | **Spearman's rho** | **Spearman's p-value** | **RMSE** | **Permutation testing p-value** |
| REST1_AP | 0.2496 | 7.30E-05 | 3.4388 | <0.0001 |
| REST1_PA | 0.1950 | 2.08E-03 | 3.4882 | 0.006 |
| REST2_AP | 0.1366 | 3.19E-02 | 3.6188 | 0.042 |
| REST2_PA | 0.1787 | 4.84E-03 | 3.5502 | 0.013 |
| CARIT | 0.2260 | 3.43E-04 | 3.4411 | 0.001 |
| FACENAME | 0.4308 | 1.38E-12 | 3.1307 | <0.0001 |
| VISMOTOR | 0.1840 | 3.71E-03 | 3.5507 | 0.009 |

**Supplementary Table 5.** Male-group predictive model performance results of the median-performing iteration of 1000 total iterations per model. All scans except those indicated by the † symbol were significant after multiple comparisons correction (based on a significance level of p=0.05; abbreviations: FN-TR, FaceName-Total Recall; RAVLT, Rey Auditory Verbal Learning Test; RAVLT-L, RAVLT-Learning; RAVLT-IR, RAVLT-Immediate Recall; RMSE, root-mean-square error).

|  | **FN-TR-predicting models** | **RAVLT-L-predicting models** | **RAVLT-IR-predicting models** |
| --- | --- | --- | --- |
| **Scan Type** | **t-test p-value** | **t-test p-value** | **t-test p-value** |
| REST1_AP | <0.0001 | <0.0001 | <0.0001 |
| REST1_PA | <0.0001 | <0.0001 | <0.0001 |
| REST2_AP | <0.0001 | <0.0001 | <0.0001 |
| REST2_PA | <0.0001 | <0.0001 | <0.0001 |
| CARIT | <0.0001 | <0.0001 | <0.0001 |
| FACENAME | <0.0001 | <0.0001 | <0.0001 |
| VISMOTOR | <0.0001 | <0.0001 | <0.0001 |

**Supplementary Table 6.** P-values from Wilcoxon rank sum tests showing that female- and male-group model performances differed across all fMRI scans for each memory score (Abbreviations: FN-TR, FaceName-Total Recall; RAVLT, Rey Auditory Verbal Learning Test; RAVLT-L, RAVLT-Learning; RAVLT-IR, RAVLT-Immediate Recall; RMSE, root-mean-square error).

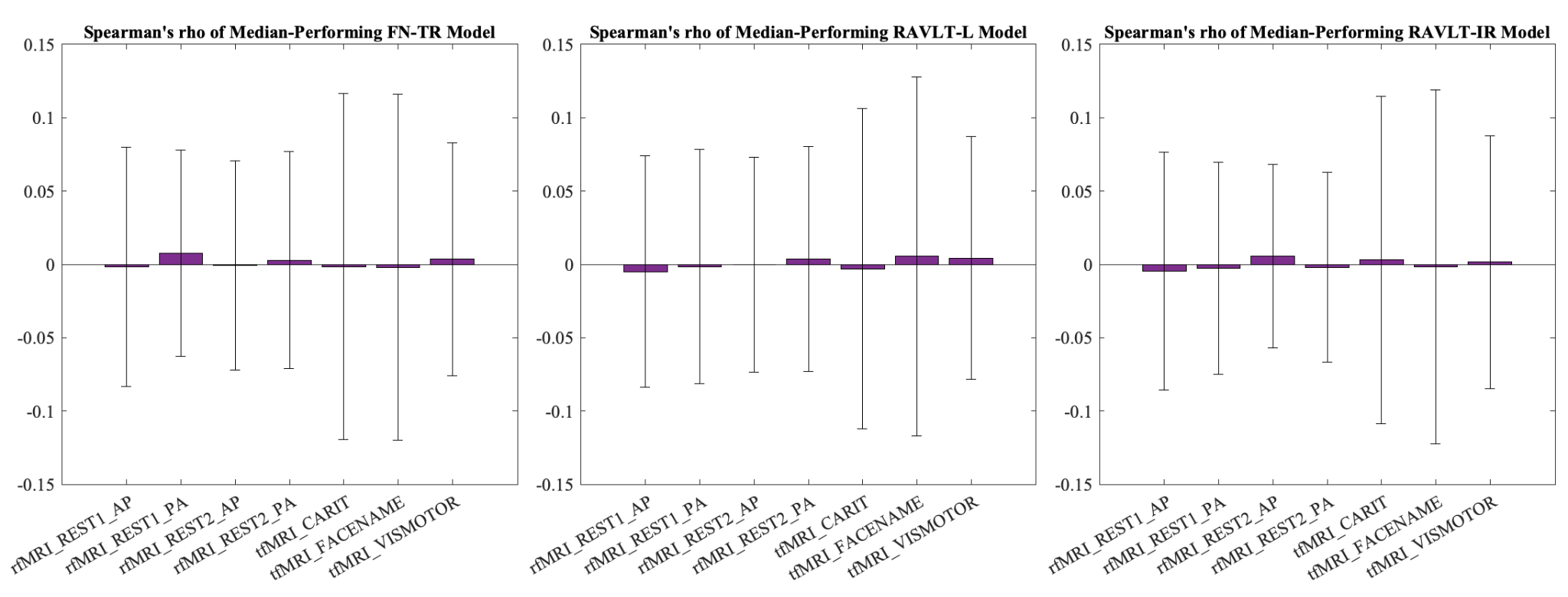

**Supplementary Figure 3.** Whole-group permutation testing (randomly shuffling participant labels prior to attempting to predict memory scores) model performance (measured using Spearman’s correlation coefficient (rho) values of the median-performing iterations of each model) across all fMRI scans for each memory metric (FN-TR, RAVLT-L, RAVLT-IR). Accuracies for all permuted models were very close to zero and significantly differed from our unpermuted models. Error bars represent standard deviations of the 1000 iterations of each model (Abbreviations: rfMRI, resting-state fMRI; tfMRI, task fMRI; FN-TR, FaceName-Total Recall; RAVLT, Rey Auditory Verbal Learning Test; RAVLT-L, RAVLT-Learning; RAVLT-IR, RAVLT-Immediate Recall).

| **FN-TR-predicting models** | | | | |
| --- | --- | --- | --- | --- |
| **Scan Type** | **Spearman's rho** | **Spearman's p-value** | **RMSE** | **Permutation testing p-value** |
| REST1_AP | -0.0018 | 0.9657 | 3.2575 | <0.0001 |
| REST1_PA | 0.0075 | 0.8588 | 3.2418 | <0.0001 |
| REST2_AP | -0.0006 | 0.9887 | 3.2562 | <0.0001 |
| REST2_PA | 0.0028 | 0.9469 | 3.2490 | <0.0001 |
| CARIT | -0.0016 | 0.9698 | 3.2500 | <0.0001 |
| FACENAME | -0.0021 | 0.9604 | 3.2561 | <0.0001 |
| VISMOTOR | 0.0035 | 0.9333 | 3.2565 | <0.0001 |
| **RAVLT-L-predicting models** | | | | |
| **Scan Type** | **Spearman's rho** | **Spearman's p-value** | **RMSE** | **Permutation testing p-value** |
| REST1_AP | -0.0049 | 0.9074 | 11.5720 | <0.0001 |
| REST1_PA | -0.0015 | 0.9722 | 11.5465 | <0.0001 |
| REST2_AP | -0.0003 | 0.9940 | 11.5683 | <0.0001 |
| REST2_PA | 0.0036 | 0.9313 | 11.5240 | <0.0001 |
| CARIT | -0.0030 | 0.9432 | 11.5973 | <0.0001 |
| FACENAME | 0.0054 | 0.8984 | 11.5674 | <0.0001 |
| VISMOTOR | 0.0043 | 0.9189 | 11.5712 | <0.0001 |
| **RAVLT-IR-predicting models** | | | | |
| **Scan Type** | **Spearman's rho** | **Spearman's p-value** | **RMSE** | **Permutation testing p-value** |
| REST1_AP | -0.0047 | 0.9114 | 3.6244 | <0.0001 |
| REST1_PA | -0.0027 | 0.9487 | 3.6172 | <0.0001 |
| REST2_AP | 0.0055 | 0.8947 | 3.6130 | <0.0001 |
| REST2_PA | -0.0021 | 0.9608 | 3.6124 | <0.0001 |
| CARIT | 0.0030 | 0.9424 | 3.6150 | <0.0001 |
| FACENAME | -0.0017 | 0.9675 | 3.6191 | <0.0001 |
| VISMOTOR | 0.0015 | 0.9719 | 3.6277 | <0.0001 |

**Supplementary Table 7.** Whole-group permutation testing (randomly shuffling participant labels prior to attempting to predict memory scores) model performance results of the median-performing iteration of 1000 total iterations per model (Abbreviations: FN-TR, FaceName-Total Recall; RAVLT, Rey Auditory Verbal Learning Test; RAVLT-L, RAVLT-Learning; RAVLT-IR, RAVLT-Immediate Recall; RMSE, root-mean-square error).

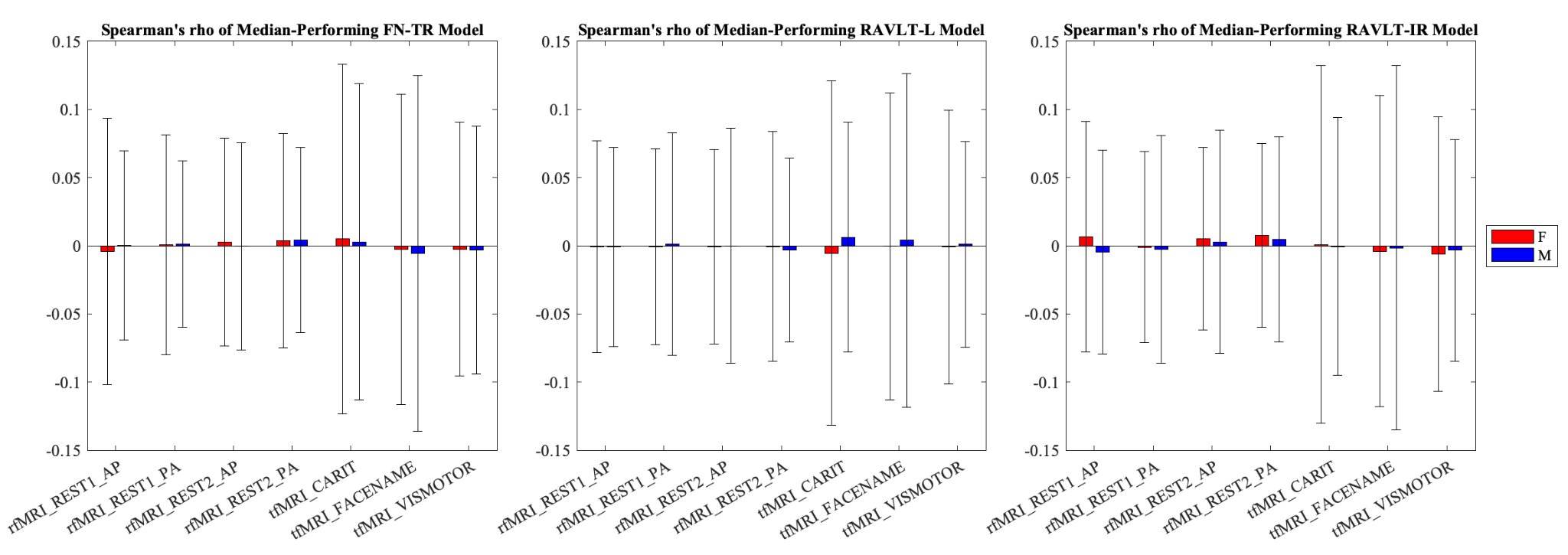

**Supplementary Figure 4.** Female- and male-group permutation testing (randomly shuffling participant labels prior to attempting to predict memory scores) model performance (measured using Spearman’s correlation coefficient (rho) values of the median-performing iterations of each model) across all fMRI scans for each memory metric (FN-TR, RAVLT-L, RAVLT-IR). Accuracies for all permuted models were very close to zero and significantly differed from our unpermuted models (with the exceptions of the three models indicated in **Supplementary Table 9**). Error bars represent standard deviations of the 1000 iterations of each model (Abbreviations: rfMRI, resting-state fMRI; tfMRI, task fMRI; FN-TR, FaceName-Total Recall; RAVLT, Rey Auditory Verbal Learning Test; RAVLT-L, RAVLT-Learning; RAVLT-IR, RAVLT-Immediate Recall).

| **FN-TR-predicting models** | | | | |
| --- | --- | --- | --- | --- |
| **Scan Type** | **Spearman's rho** | **Spearman's p-value** | **RMSE** | **Permutation testing p-value** |
| REST1_AP | -0.0042 | 0.9404 | 3.0321 | <0.0001 |
| REST1_PA | 0.0007 | 0.9902 | 3.0344 | <0.0001 |
| REST2_AP | 0.0027 | 0.9621 | 3.0219 | 0.001 |
| REST2_PA | 0.0036 | 0.9495 | 3.0297 | 0.001 |
| CARIT | 0.0050 | 0.9296 | 2.9932 | <0.0001 |
| FACENAME | -0.0028 | 0.9598 | 3.0269 | <0.0001 |
| VISMOTOR | -0.0026 | 0.9625 | 3.0036 | <0.0001 |
| **RAVLT-L-predicting models** | | | | |
| **Scan Type** | **Spearman's rho** | **Spearman's p-value** | **RMSE** | **Permutation testing p-value** |
| REST1_AP | -0.0007 | 0.9905 | 11.7344 | <0.0001 |
| REST1_PA | -0.0007 | 0.9897 | 11.7603 | 0.023 |
| REST2_AP | -0.0009 | 0.9869 | 11.7388 | <0.0001 |
| REST2_PA | -0.0006 | 0.9917 | 11.7487 | <0.0001 |
| CARIT | -0.0054 | 0.9233 | 11.6726 | <0.0001 |
| FACENAME | -0.0005 | 0.9934 | 11.7154 | <0.0001 |
| VISMOTOR | -0.0009 | 0.9873 | 11.6628 | <0.0001 |
| **RAVLT-IR-predicting models** | | | | |
| **Scan Type** | **Spearman's rho** | **Spearman's p-value** | **RMSE** | **Permutation testing p-value** |
| REST1_AP | 0.0067 | 0.9037 | 3.4037 | <0.0001 |
| REST1_PA | -0.0010 | 0.9856 | 3.4135 | <0.0001 |
| REST2_AP | 0.0052 | 0.9249 | 3.3953 | <0.0001 |
| REST2_PA | 0.0075 | 0.8925 | 3.4075 | <0.0001 |
| CARIT | 0.0010 | 0.9860 | 3.3843 | <0.0001 |
| FACENAME | -0.0040 | 0.9423 | 3.4036 | <0.0001 |
| VISMOTOR | -0.0063 | 0.9098 | 3.3956 | <0.0001 |

**Supplementary Table 8.** Female-group permutation testing (randomly shuffling participant labels prior to attempting to predict memory scores) model performance results of the median-performing iteration of 1000 total iterations per model (Abbreviations: FN-TR, FaceName-Total Recall; RAVLT, Rey Auditory Verbal Learning Test; RAVLT-L, RAVLT-Learning; RAVLT-IR, RAVLT-Immediate Recall; RMSE, root-mean-square error).

| **FN-TR-predicting models** | | | | |
| --- | --- | --- | --- | --- |
| **Scan Type** | **Spearman's rho** | **Spearman's p-value** | **RMSE** | **Permutation testing p-value** |
| REST1_AP | 0.0002 | 0.9970 | 3.3656 | 0.061 † |
| REST1_PA | 0.0011 | 0.9867 | 3.3649 | 0.457 † |
| REST2_AP | -0.0005 | 0.9944 | 3.3692 | 0.03 |
| REST2_PA | 0.0041 | 0.9499 | 3.3675 | 0.003 |
| CARIT | 0.0028 | 0.9662 | 3.3195 | <0.0001 |
| FACENAME | -0.0056 | 0.9316 | 3.3366 | <0.0001 |
| VISMOTOR | -0.0033 | 0.9598 | 3.3484 | 0.004 |
| **RAVLT-L-predicting models** | | | | |
| **Scan Type** | **Spearman's rho** | **Spearman's p-value** | **RMSE** | **Permutation testing p-value** |
| REST1_AP | -0.0008 | 0.9905 | 12.5513 | 0.006 |
| REST1_PA | 0.0013 | 0.9834 | 12.5541 | 0.006 |
| REST2_AP | -0.0001 | 0.9992 | 12.5799 | 0.025 |
| REST2_PA | -0.0033 | 0.9589 | 12.5775 | 0.027 |
| CARIT | 0.0063 | 0.9218 | 12.4378 | 0.001 |
| FACENAME | 0.0041 | 0.9499 | 12.4685 | <0.0001 |
| VISMOTOR | 0.0010 | 0.9870 | 12.5259 | 0.053 † |
| **RAVLT-IR-predicting models** | | | | |
| **Scan Type** | **Spearman's rho** | **Spearman's p-value** | **RMSE** | **Permutation testing p-value** |
| REST1_AP | -0.0048 | 0.9397 | 3.7553 | <0.0001 |
| REST1_PA | -0.0027 | 0.9661 | 3.7484 | 0.006 |
| REST2_AP | 0.0029 | 0.9641 | 3.7458 | 0.042 |
| REST2_PA | 0.0045 | 0.9438 | 3.7441 | 0.013 |
| CARIT | -0.0005 | 0.9934 | 3.7048 | 0.001 |
| FACENAME | -0.0015 | 0.9811 | 3.7165 | <0.0001 |
| VISMOTOR | -0.0034 | 0.9582 | 3.7309 | 0.009 |

**Supplementary Table 9.** Male-group permutation testing (randomly shuffling participant labels prior to attempting to predict memory scores) model performance results of the median-performing iteration of 1000 total iterations per model. † indicates the models that did not survive correction for multiple comparisons (based on a significance level of p=0.05; abbreviations: FN-TR, FaceName-Total Recall; RAVLT, Rey Auditory Verbal Learning Test; RAVLT-L, RAVLT-Learning; RAVLT-IR, RAVLT-Immediate Recall; RMSE, root-mean-square error).

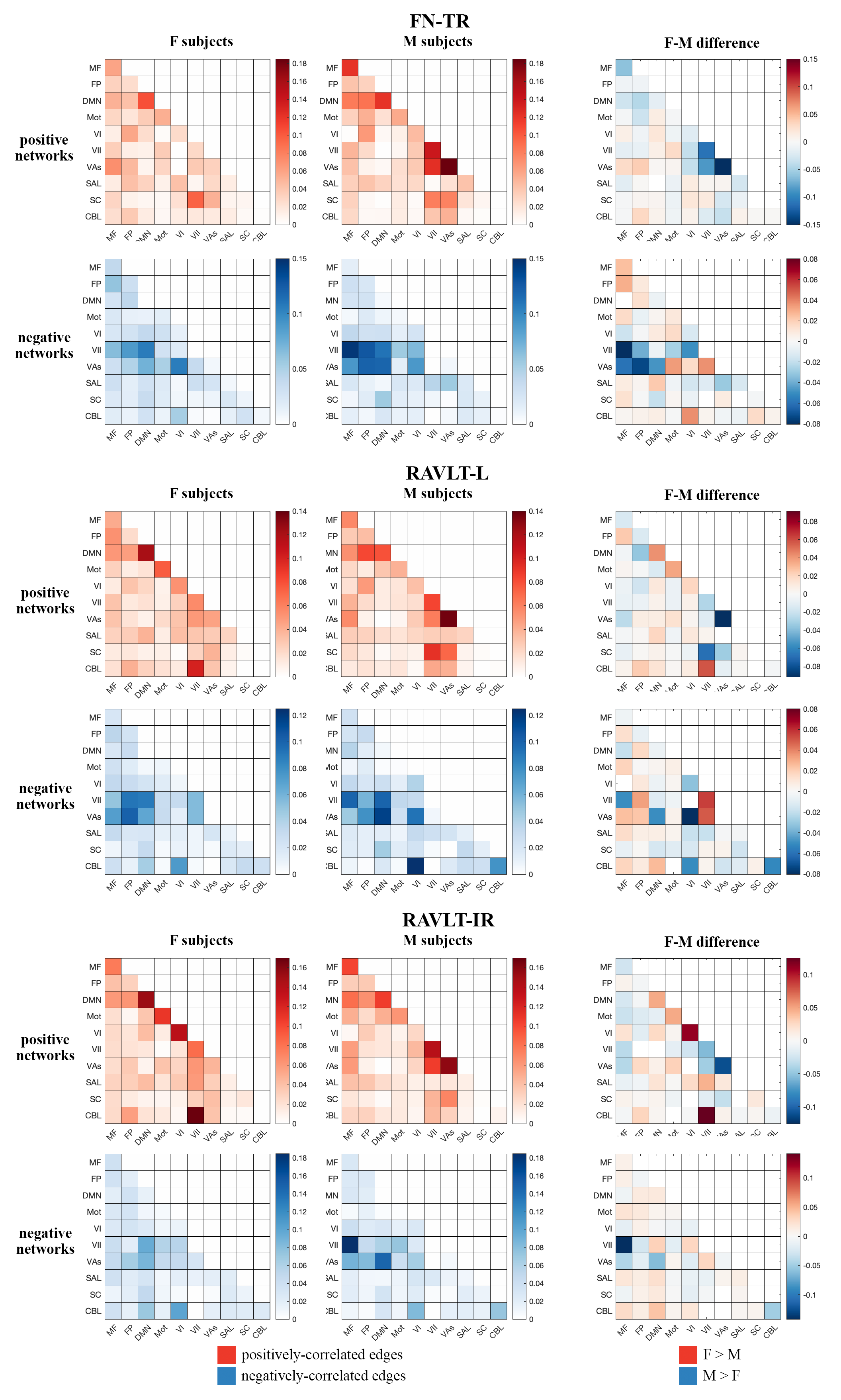

**Supplementary Figure 5.** Inter-network edge heatmaps for each sex separately and the difference between the sexes (male-group edges subtracted from female-group edges) based on the 10-functional-network parcellation of the Shen 268 node atlas. We observed sex differences in key edges across all FaceName tfMRI models, particularly in intra-DMN and intra-visual (VI, VII, and VAs)-network edges, as well as edges between the MF, DMN, and visual networks (Abbreviations: F, female; M, male; MF, medial frontal; FP, fronto-parietal; DMN, default mode network; Mot, motor; VI, visual I; VII, visual II; VAs, visual association areas; SAL, limbic; SC, basal ganglia; CBL, cerebellum; FN-TR, FaceName-Total Recall; RAVLT, Rey Auditory Verbal Learning Test; RAVLT-L, RAVLT-Learning; RAVLT-IR, RAVLT-Immediate Recall).

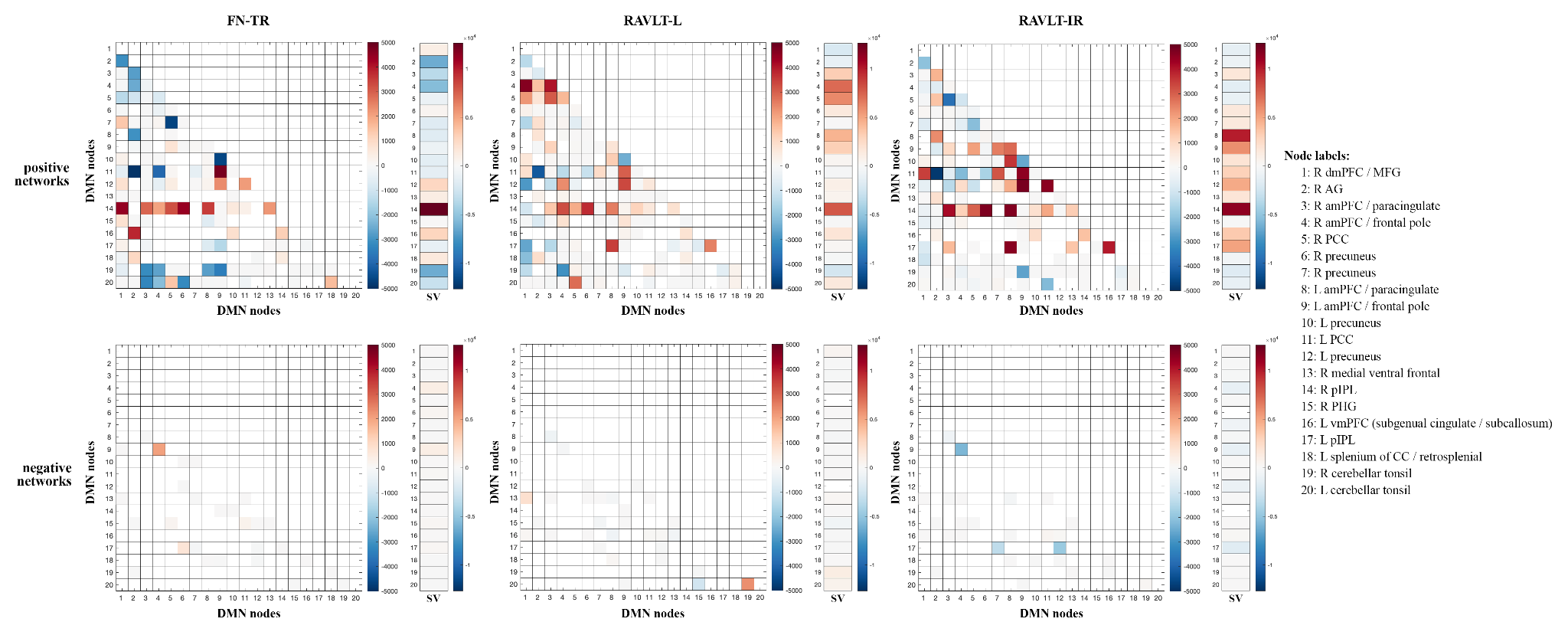

**Supplementary Figure 6.** Intra-DMN edge heatmap for RAVLT-IR, RAVLT-L, and FN-TR models. Positively-associated (top row) and negatively-associated (bottom row) edges that are enriched in the three models evaluated. Values of each edge were calculated by subtracting female and male edge numbers (F-M) from the model (Abbreviations: RAVLT-IR, RAVLT-Immediate Recall; L, left; R, right; dmPFC, dorsomedial prefrontal cortex; MFG, middle frontal gyrus; AG, angular gyrus; aMPFC, anterior medial prefrontal cortex; PCC, posterior cingulate cortex; pIPL, posterior inferior parietal lobe; PHG, parahippocampal gyrus; vmPFC, ventromedial prefrontal cortex; CC, corpus collosum; SV, summed vector).
